# Piperidine CD4-mimetic compounds expose vulnerable Env epitopes sensitizing HIV-1-infected cells to ADCC

**DOI:** 10.1101/2023.03.23.533923

**Authors:** Shilei Ding, William D. Tolbert, Huile Zhu, Daniel Lee, Tyler Higgins, Xuchen Zhao, Dung Nguyen, Rebekah Sherburn, Jonathan Richard, Gabrielle-Gendron Lepage, Halima Medjahed, Mohammadjavad Mohammadi, Cameron Abrams, Marzena Pazgier, Amos B. Smith, Andrés Finzi

## Abstract

The ability of HIV-1 accessory proteins Nef and Vpu to decrease CD4 levels contributes to the protection of infected cells from antibody-dependent cellular cytotoxicity (ADCC) by preventing the exposure of Env vulnerable epitopes. Small-molecule CD4 mimetics (CD4mc) based on the indane and piperidine scaffolds such as (+)-BNM-III-170 and (*S*)-MCG-IV-210 sensitize HIV-1 infected cells to ADCC by exposing CD4-induced (CD4i) epitopes recognized by non-neutralizing antibodies abundantly present in plasma from people living with HIV. Here, we characterize a new family of CD4mc, (*S*)-MCG-IV-210 derivatives, based on the piperidine scaffold which engage the gp120 within the Phe43 cavity by targeting the highly-conserved Asp^368^ Env residue. We utilized structure-based approaches and developed a series of piperidine analogs with improved activity to inhibit infection of difficult-to-neutralize tier-2 viruses and sensitize infected cells to ADCC mediated by HIV+ plasma. Moreover, the new analogs formed an H-bond with the α-carboxylic acid group of Asp^368^, opening a new avenue to enlarge the breadth of this family of anti-Env small molecules. Overall, the new structural and biological attributes of these molecules make them good candidates for strategies aimed at the elimination HIV-1-infected cells.

## 1. Introduction

HIV-1 envelope glycoproteins (Env) mediate virus infection by binding to CD4 on the surface of host cells [1-3]. Env is constituted of a trimer of heterodimers made of a transmembrane (gp41) and surface (gp120) subunit. Upon CD4 binding, the gp120 ensues a series of conformational changes leading to the exposure of the coreceptor binding site (CoRBS) and the gp41 helical heptad repeat (HR1) [4]. These CD4-induced (CD4i) epitopes can be recognized by non-neutralizing antibodies (nnAbs) abundantly present in plasma of people living with HIV (PLWH) [5], some of which are able to mediate antibody-dependent cellular cytotoxicity (ADCC) [6, 7].

Small-molecule CD4-mimetic compounds (CD4mc) trigger similar conformational changes as CD4 by engaging the gp120 within the highly conserved region of Env that accommodates Phe43 of CD4 (referred to as the Phe43 cavity) [8-11]. Based on the original compounds NBD-556 and NBD-557 discovered by Debnath [12], derivatives such as JP-III-48 or (+)-BNM-III-170 were developed with better capacity to neutralize HIV-1 virus particles and sensitize HIV-1 infected cells to ADCC [9, 10, 13-17]. Further screening of small molecules led to the finding of piperidine analog (*S*)-MCG-IV-210, which engages the Phe43 cavity in a similar manner to that of (+)-BNM-III-170 while being in closer proximity to the highly conserved CD4-binding residue Asp^368^ [18].

In this study, analogs of the piperidine (*S*)-MCG-IV-210 were designed and synthesized with the goal to improve their interaction with Asp^368^. The analogs development was guided by high resolution structures of the complexes formed by newly-developed CD4mc and a gp120 Env core. The structures provide insights into analogs’ interactions within the CD4-binding cavity and how they specifically interact with Asp^368^. The capacity of this new series of compounds to neutralize viral particles and sensitize infected cells to ADCC was measured.

## 2. Results

### 2.1. New piperidine analogs

Recently, we reported on the identification and development of a new class of piperidine-based small molecules, namely (*S*)-MCG-IV-210 and derivatives thereof, exhibiting anti-viral properties that lead to sensitization of HIV-1 infected cells to ADCC [18, 19]. The structure of (*S*)-MCG-IV-210 features a pendant 4-chloro-3-fluoro-arene (Region I) which inserts deeply into the Phe43 cavity, an amide linker (Region II) that extends the molecule out of the Phe43 cavity, and a piperidine ring (Region III) that has been determined to be essential for the anti-viral activity (Figure 1A).

**Figure 1.**
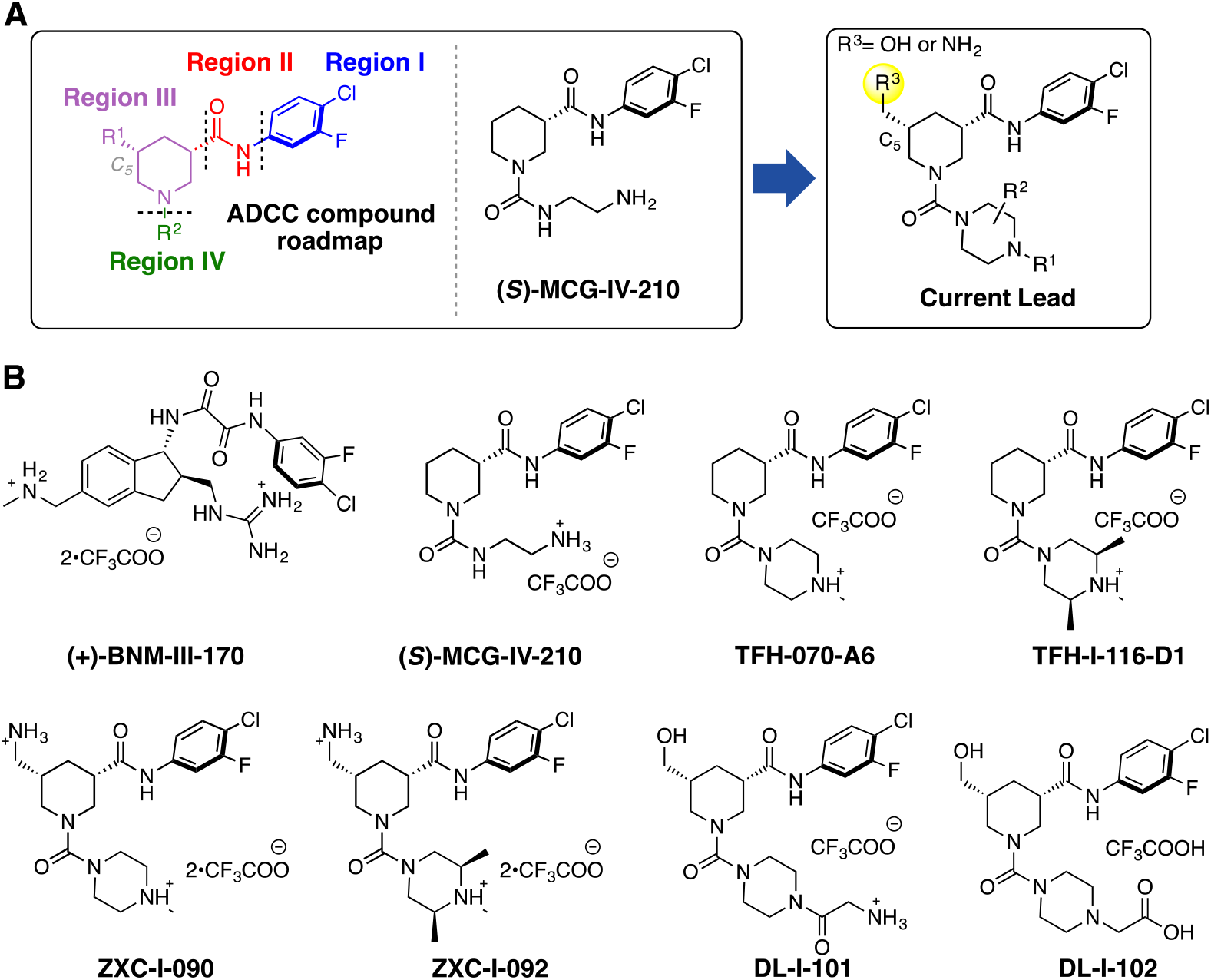
(**A**) A structural roadmap (i.e., Region I-IV) of the lead molecule scaffold. (**B**) Chemical structures of (+)-BNM-III-170, (S)-MCG-IV-210, TFH-070-A6, TFH-I-116-D1, ZXC-I-090, ZXC-I-092, DL-I-101, and DL-I-102.

In this study, we present new analogs where the linear urea in Region IV of the molecule skeleton has been replaced with heterocyclic groups such as a methyl-substituted piperazine (TFH-070-A6, Figure 1B) or a functionalized piperazine (TFH-I-116-D1, Figure 1B). To enable the molecule to contact with the Asp^368^ carboxylate side chain, we modified the C_5_ position of the piperidine by adding a hydrogen bonding promoter such as *i)* primary amines on TFH-070-A6 or TFH-I-116-D1 to make ZXC-I-090 or ZXC-I-092 (Figure 1B) and *ii)* hydroxy methyl groups found in DL-I-101 or DL-I-102, with further modifications to the Region IV based on the previously reported (*S*)-MCG-IV-210 scaffold.

### 2.2. The Design and Synthesis of New CD4mc Piperidine Analogs

We began the structure-based design of the new CD4mc piperidine analogs by comparing high-resolution crystal structures of complexes of clade A/E gp120_CRF01_ LMHT core with (*S*)-MCG-IV-210 and indane CD4mc (+)-BNM-III-170 (Figure 2). While both small molecules CD4mcs contacted Gly^472^, we observed in the gp120 complex of (*S*)-MCG-IV-210 that the linear urea in region IV of the molecule did not engage the β20-21 loop (Figure 2C). However, we observed that the C_5_ position of the piperidine scaffold was in close proximity to the highly conserved Asp^368^ side chain residue within the Phe43 cavity on the gp_120_ complex.

**Figure 2.**
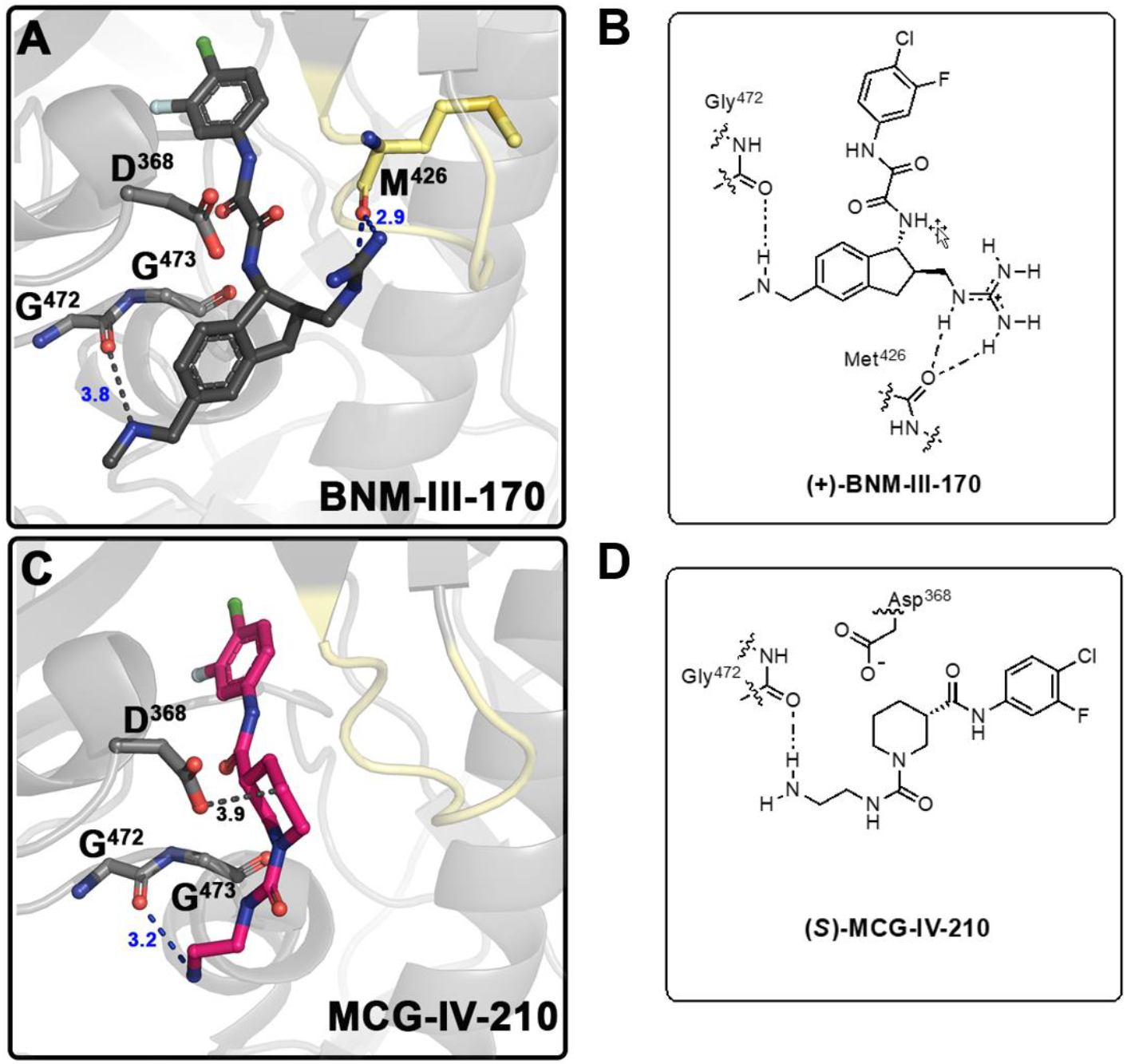
Development of CD4mc Piperidine Analogs. Structure of the complex of (+)-BNM-III-170 (PDB code: 6UT1) (**A**) and (S)-MCG-IV-210 (PDB code: 6USW) (**C**) bound to clade A/E gp120_CRF01_ LMHT core [10, 18]. H-bonds are shown as blue dashed lines and other contacts to Gly^472^ Met ^426^and Asp^368^ as grey dashed lines. (**B**) Graphical illustration of (+)-BNM-III-170 in contact with Gly^472^ and Met^426^. (**D**) Graphical illustration of (S)-MCG-IV-210 in contact with Gly^472^.

To this end, we initiated our study by the derivatizations of various regions of the (*S*)-MCG-IV-210 scaffold with the aim of *i)* continuing the structure optimization of Region IV to probe the β20-21 loop within the gp120 complex and *ii)* incorporating a polar moiety on C_5_ of the piperidine ring (Region III) to establish an electrostatic interaction with Asp^368^.

### 2.3. Biological Testing

We first tested the ability of the new piperidine analogs to expose the co-receptor binding site (CoRBS) by measuring binding of the anti-CoRBS 17b mAb to infected cells. Briefly, we infected primary CD4+ T cells with HIV-1_CH58TF_, HIV-1_JRFL_ or HIV-1_AD8_. Two days later, infected cells were co-incubated with 50 µM (+)-BNM-III-170, (*S*)-MCG-IV-210, TFH-070-A6, TFH-I-116-D1, ZXC-I-090, ZXC-I-092, DL-I-101, and DL-I-102 or the same volume of DMSO, and the 17b interaction was measured by flow cytometry after intracellular anti-p24 to identify the infected cell population. As previously reported [18], (*S*)-MCG-IV-210 exposed the CoRBS of cells infected with HIV-1_CH58_, but not the tier-2 HIV-1_JRFL_ (Figure 3A, 3B). Interestingly, the newly developed analogs -TFH-070-A6, TFH-I-116-D1, ZXC-I-092, DL-I-101, and DL-I-102 successfully exposed the CoRBS of HIV-1_JRFL_ infected cells, albeit to a lower level than the potent (+)-BNM-III-170 (Figure 3B). A similar phenotype was observed when using HIV-1_AD8_ infected cells (Figure 3C). This confirms the superior activity of the new analogs over the previous lead for this new class of CD4mc.

**Figure 3.**
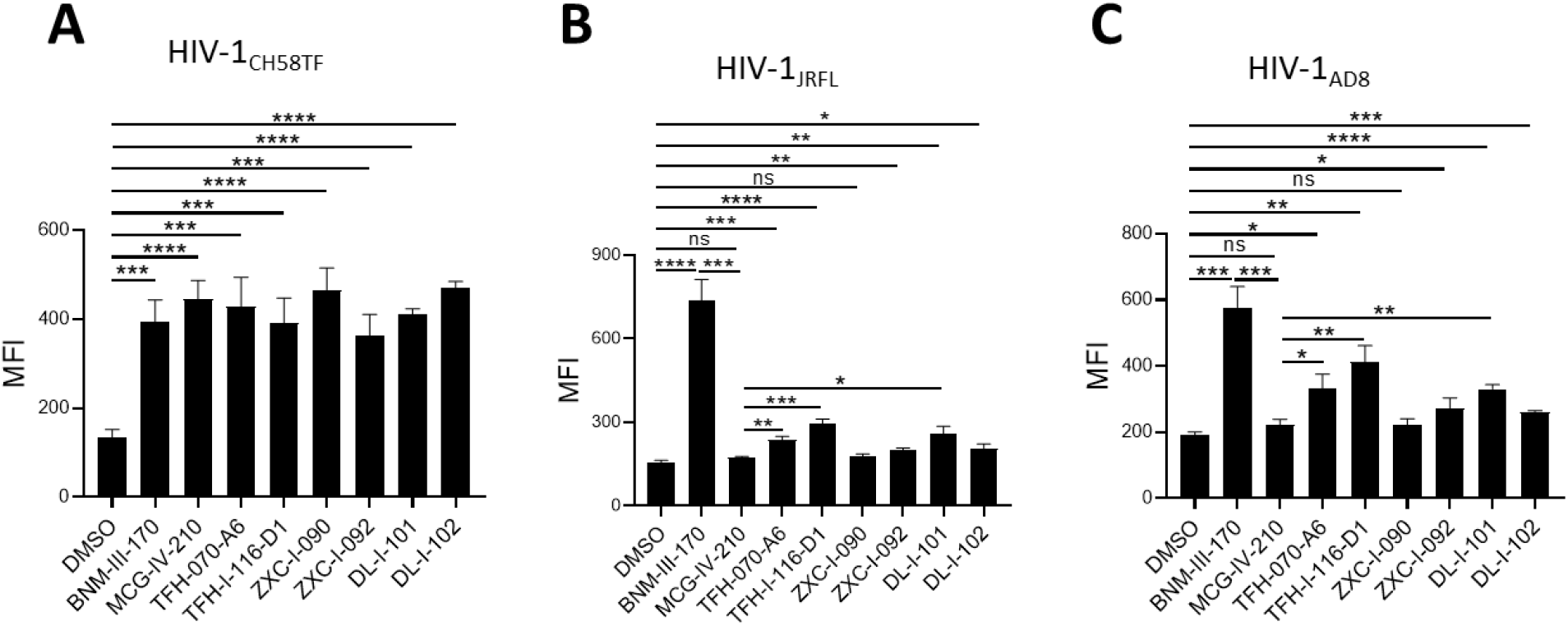
Anti-CoRBS Ab binding. Anti-CoRBS Ab – 17b binding to (**A**) HIV-1_CH58TF_, (**B**) HIV-1_JRFL_ (**C**) or HIV-1_AD8_ infected primary CD4+ T cells was done in the presence of 50 µM of the indicated CD4mcs or the same volume of vehicle (DMSO). The Median fluorescence intensity (MFI) of 17b binding is reported for the productively-infected cell (p24+) population. Data shown are the mean ± SD of at least three independent experiments. Statistical significance was evaluated using an unpaired *t* test (*, P < 0.05; **, P < 0.01; ***, P < 0.001; ****, P < 0.0001; ns, not significant).

### 2.4. Structural basis of interaction of (S)-MCG-IV-210 derivatives with a gp120 core

To determine molecular details of the interaction of the new piperidine analogs with Env we solved high resolution crystal structures of TFH-070-A6, TFH-I-116-D1, ZXC-I-090, ZXC-I-092, DL-I-101, and DL-I-102 bound to LMHT gp120_CRF01_AE_ core_e_ [10, 18]. Structures were solved from the high resolution of 2.2 Å (DL-I-101) to the lower resolution of 3.5 Å (TFH-I-116-D1). Data collection and refinement statistics are shown in Table 1. In Figure 4 the piperidine analogs are shown within the CD4 (Phe43) binding cavity with (+)-BNM-III-170 and (*S*)-MCG-IV-210 superimposed to highlight differences in their binding modes. All analogs utilize the 4-chloro-3-fluoro-arene and amide linker to anchor deeply into the Phe43 cavity with largely identical positions and orientations within the cavity (Figure 4B and C). Major differences, however, are observed for the conformations of the piperidine scaffold with the new compounds binding in close proximity to the gp120 β20-21 loop region while preserving contacts to the highly conserved Asp^368^ side chain.

**Table 1.** Data collection and refinement statistics.

|  | LM/HT<br>gp120<sub>CRF01_AE</sub><br>core<sub>e</sub> – TFH-I-070-A6 | LM/HT<br>gp120<sub>CRF01_AE</sub><br>core<sub>e</sub> – TFH-I-116-D1 | LM/HT<br>gp120<sub>CRF01_AE</sub><br>core<sub>e</sub> – ZXC-I-090 | LM/HT<br>gp120<sub>CRF01_AE</sub><br>core<sub>e</sub> – ZXC-I-092 | LM/HS<br>gp120<sub>CRF01_AE</sub><br>core<sub>e</sub> – DL-I-101 | LM/HS<br>gp120<sub>CRF01_AE</sub><br>core<sub>e</sub> – DL-I-102 |
| --- | --- | --- | --- | --- | --- | --- |
| **Data collection** |  |  |  |  |  |  |
| Wavelength, Å | 0.979 | 0.979 | 0.979 | 0.979 | 0.979 | 0.979 |
| Space group | P2<sub>1</sub>2<sub>1</sub>2<sub>1</sub> | P2<sub>1</sub>2<sub>1</sub>2<sub>1</sub> | P2<sub>1</sub>2<sub>1</sub>2<sub>1</sub> | P2<sub>1</sub>2<sub>1</sub>2<sub>1</sub> | P2<sub>1</sub>2<sub>1</sub>2<sub>1</sub> | P2<sub>1</sub>2<sub>1</sub>2<sub>1</sub> |
| Cell parameters<br>a, b, c, Å | 63.6, 66.3,<br>91.4 | 64.7, 66.0,<br>87.8 | 61.8, 65.9,<br>91.2 | 60.0, 65.2,<br>87.3 | 67.7, 66.9,<br>92.2 | 194.5, 86.6,<br>57.7 |
| α, β, γ, ° | 90, 90, 90 | 90, 90, 90 | 90, 90, 90 | 90, 909, 90 | 90, 90, 90 | 90, 90, 90 |
| Molecules/a.u. | 1 | 1 | 1 | 1 | 1 | 1 |
| Resolution, (Å) | 50-2.4 (2.44-2.4) | 50-3.5 (3.69-3.5) | 50-2.4 (2.44-2.4) | 50-2.85 (2.9-2.85) | 50-2.19<br>(2.23-2.19) | 50-2.6 (2.64-2.6) |
| # of reflections |  |  |  |  |  |  |
| Total | 49,666 | 11,650 | 61,106 | 34,488 | 86,920 | 39,768 |
| Unique | 13,796 | 4,021 | 14,904 | 7,185 | 20,214 | 10,748 |
| R<sub>merge</sub><sup>a</sup>, % | 9.1 (57.9) | 22.3 (113) | 10.6 (80.1) | 12.1 (104) | 17.2 (66.5) | 15.4 (98.3) |
| R<sub>pim</sub><sup>b</sup>, % | 5.1 (32.3) | 14.7 (72.9) | 5.8 (71.1) | 8.2 (77.4) | 9.7 (36.4) | 9.0 (65.0) |
| CC<sub>1/2</sub><sup>c</sup> | 0.98 (0.71) | 0.97 (0.33) | 1.0 (0.50) | 1.0 (0.42) | 1.0 (0.65) | 0.94 (0.60) |
| I/σ | 12.3 (1.3) | 2.8 (0.6) | 14.2 (0.8) | 12.5 (0.8) | 25.5 (2.1) | 17.3 (1.1) |
| Completeness,<br>% | 88.3 (78.2) | 80.6 (83.9) | 97.8 (86.4) | 83.3 (78.0) | 97.4 (85.0) | 86.1 (90.2) |
| Redundancy | 3.6 (3.7) | 2.9 (3.0) | 4.1 (2.7) | 4.8 (4.2) | 4.3 (3.8) | 3.7 (3.1) |
| **Refinement Statistics** |  |  |  |  |  |  |
| Resolution, Å | 50.0 – 2.4 | 50.0 – 3.5 | 50.0 – 2.4 | 50.0 – 2.85 | 50.0 – 2.19 | 50.0 – 2.6 |
| R<sup>d</sup> % | 22.1 | 25.6 | 24.0 | 24.2 | 20.8 | 22.1 |
| R<sub>free</sub><sup>e</sup>, % | 27.4 | 30.7 | 26.7 | 29.6 | 25.5 | 27.4 |
| # of atoms |  |  |  |  |  |  |
| Protein | 2,668 | 2,640 | 2,665 | 2,675 | 2,683 | 2,668 |
| Water | 32 | – | 23 | – | 64 | 2 |
| Ligand/Ion | 212 | 184 | 211 | 186 | 193 | 186 |
| Overall B value<br>(Å)<sup>2</sup> |  |  |  |  |  |  |
| Protein | 52 | 82 | 62 | 73 | 53 | 85 |
| Water | 48 | – | 56 | – | 49 | 74 |
| Ligand/Ion | 70 | 96 | 74 | 86 | 63 | 102 |
| RMSD<sup>f</sup> |  |  |  |  |  |  |
| Bond lengths, Å | 0.005 | 0.004 | 0.006 | 0.009 | 0.007 | 0.012 |
| Bond angles, ° | 1.0 | 0.79 | 0.98 | 1.3 | 1.0 | 1.4 |
| Ramachandran<sup>g</sup> |  |  |  |  |  |  |
| favored, % | 94.0 | 90.2 | 94.0 | 89.5 | 95.0 | 92.5 |
| allowed, % | 4.2 | 7.4 | 5.4 | 9.0 | 4.7 | 7.2 |
| outliers, % | 1.8 | 2.4 | 0.6 | 1.5 | 0.3 | 0.3 |
| PDB ID | 8GD0 | 8GJT | 8GCZ | 8GD1 | 8GD3 | 8GD5 |

Values in parentheses are for highest-resolution shell
<sup>a</sup>R<sub>merge</sub> = $\sum |I - \langle I \rangle| / \sum I$ , where $I$ is the observed intensity and $\langle I \rangle$ is the average intensity obtained from multiple observations of symmetry-related reflections after rejections
<sup>b</sup>R<sub>pim</sub> = as defined in [20]
<sup>c</sup>CC<sub>1/2</sub> = as defined by Karplus and Diederichs [21]
<sup>d</sup>R = $\sum \|F_o| - |F_c\| / \sum |F_o|$ , where $F_o$ and $F_c$ are the observed and calculated structure factors, respectively
<sup>e</sup>R<sub>free</sub> = as defined by Brünger [22]
<sup>f</sup>RMSD = Root mean square deviation
<sup>g</sup>Calculated with MolProbity

**Figure 4.**
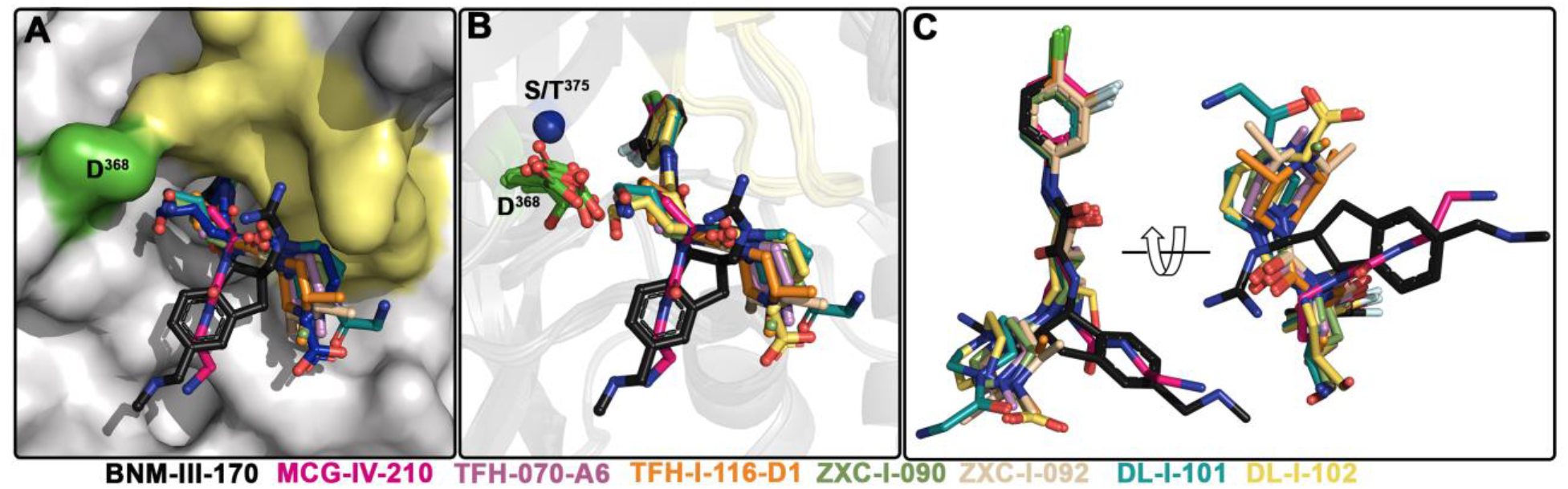
Crystal structures of (+)-BNM-III-170, TFH-070-A6, TFH-I-116-D-1,TFH-II-151, ZXC-I-090, ZXC-I-092, DL-I-101, and DL-I-102 derivatives in complex with a gp120_CRF01_AE_ LMHS or LMHT core_e_. (**A**) and (**B**) (+)-BNM-III-170, (*S*)-MCG-IV-210, and piperidine analogs within the Phe^43^ cavity (**A**) gp120 is shown as a surface with the B20-21 loop region and D^368^ colored pale yellow and green, respectively. (**B**) gp120 is show as ribbon with the B21-21 loop region colored yellow, and D^368^ and Ser/Thr^375^ are shown as green sticks or as a blue sphere. (**C**) Superimposition of the piperidine derivatives onto (+)-BNM-III-170 and (*S*)-MCG-IV-210.

Figure 5 shows the details of the interaction of the new compounds within the CD4 binding pocket and the contact network to Asp^368^, Glu^370^ (Figure 5A), and the β20-21 loop region (Figure 5C). A direct comparison of the total buried surface area (BSA) of each compound-gp120 core_e_ complex (Figure 5B) indicates that there is a significant increase in the buried surface for the new compounds as compared to their prototype (*S*)-MCG-IV-210. The highest BSAs are observed for ZXC-I-092 and DL-I-102 with 843.2 Å^2^ and 826.0 Å^2^, respectively (as compared to 699.1 Å^2^ for (*S*)-MCG-IV-210). Analyses of the BSA of individual gp120 residues that contribute to the compound interface (Figure 5B) confirm that the new analogs greatly capitalize upon interactions with Asp^368^, Glu^370^, and the β20-21 gp120 loop. A few of the compounds (ZXC-I-092, DL-I-101 and DL-I-102) also reach Arg^476^. The latter represents a new contact region that has not previously been targeted by this class of CD4mc compounds.

**Figure 5.**
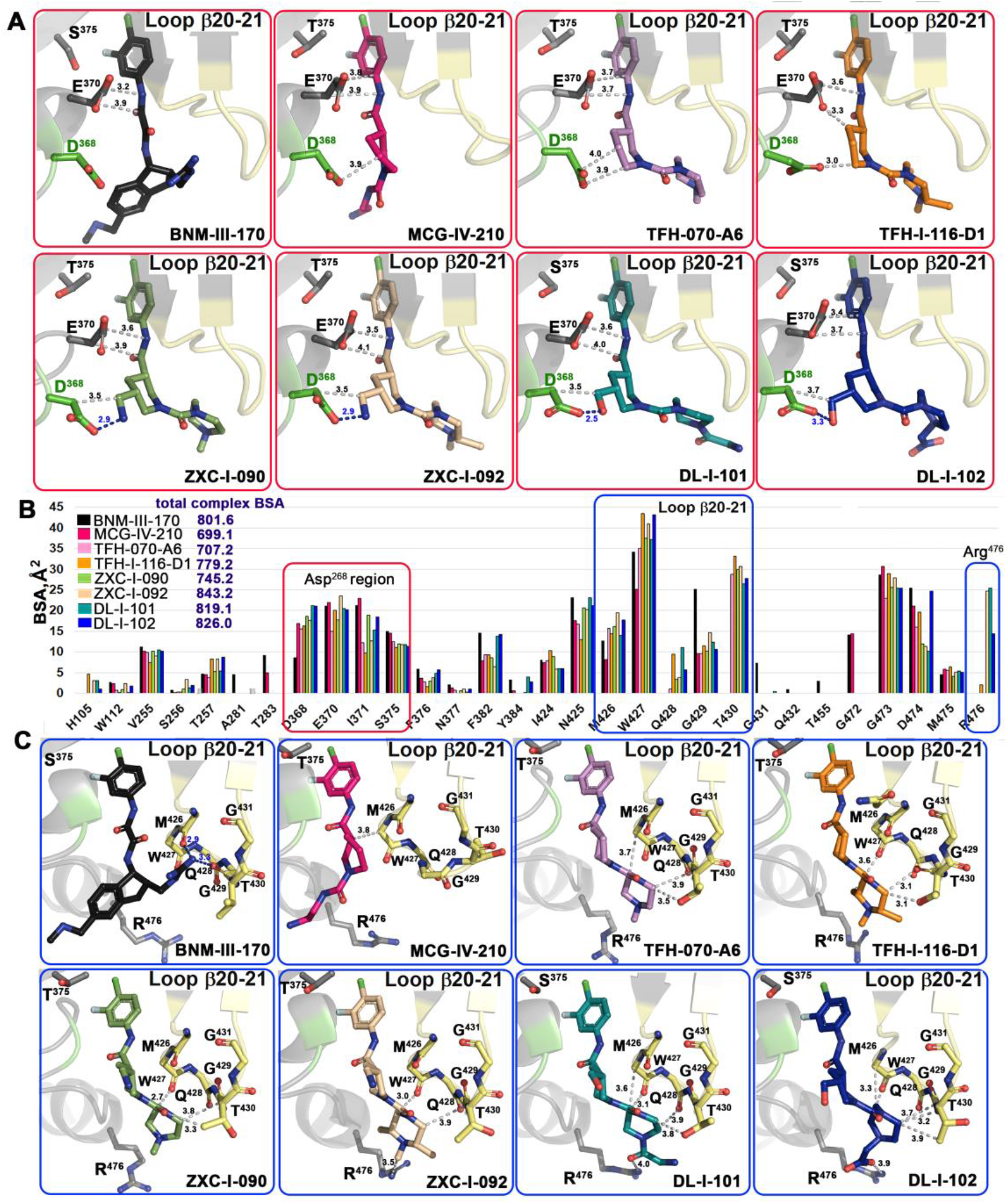
Molecular details of the interaction of piperidine analogs with the Phe43 cavity of HIV-1 Env. (**A**) and (**C**) details of each compound’s interaction with Asp^368^ and β20-21 loop region, respectively. Crystal structures of each analog in complex with gp120_CRF01_AE_ core_e_ were superimposed based upon gp120 and identical views shown with H-bonds as blue dashed lines. The distances to Asp^368^, Glu^370^, the main chain atoms of the β20-21 loop (residues from Met^426^ to Gly^431^), and the side chains of Thr^430^ and Arg^476^ are shown as grey dashed lines with distances in Å. (**B**) The relative contribution of individual gp120 residues to the binding interface for each compound shown as a buried surface area (BSA) value as calculated by PISA. The buried surface area represents the solvent-accessible surface area of the corresponding residue that is buried upon interface formation. The total BSA for each complex is shown next to the compound label.

One of the major goals for the structure-based development of this class of piperidine analogs was to make contact to the side chains of Asp^368^ and Glu^370^, among the most conserved Phe43 cavity lining residues among HIV-1 clades. Four of the new analogs achieve this goal (ZXC-I-090, ZXC-I-092, DL-I-101, and DL-I-102) by placing an amine nitrogen or hydroxy methyl oxygen of the piperazine ring at a hydrogen bond distance to the carboxyl group of Asp^368^. ZXC-I-090 and ZXC-I-092 both use their amines to establish 2.9 Å salt bridges with Asp^368^ while DL-I-101 and DL-I-102 use the hydroxyl of their hydroxy methyl to make 2.5 Å and 3.3 Å H-bonds with Asp^368^, respectively. All of the analogs also simultaneously bind close to Glu^370^ with distances to side chain atoms in the range of 3.2-4.1Å.

New for this class of analogs is the ability to rely on binding to the main chain atoms of the β20-21 loop (gp120 residues 426 to 430) (Figure 5C). These include an intensive network of interactions to the main chain oxygens of Met^426^ and Trp^427^, the backbone of Gln^428^, and the side chain of Thr^431^. The BSA for individual residues in this region is significantly higher than equivalent residues buried at the interfaces of (+)-BNM-III-170 and the prototype (*S*)-MCG-IV-210. Two compounds (DL-I-101 and DL-I-102) add to this network with contacts to Arg^476^ (Figure 5C).

### 2.5. (S)-MCG-IV-210 derivatives neutralize HIV-1 viral particles

We next evaluated the capacity of (*S*)-MCG-IV-210 derivatives to neutralize viral particles of HIV-1_CH58TF_, HIV-1_JRFL_ or HIV-1_AD8_ by using a standard TZM-bl neutralization assay. As a positive control, we used (+)-BNM-III-170 [23]. A VSV-G pseudovirus was used as a negative control. All tested derivatives were specific to HIV-1 Env since no effect was observed with VSV-G pseudoviruses (Figure 6A). As expected, all tested derivatives neutralized HIV-1_CH58TF_ at low micromolar concentrations (Figure 6A), especially TFH-I-116-D1 with an IC_50_ of 0.06548 µM, which were close to that of (+)-BNM-III-170 (0.03358 µM) (Figure 6A, Table 2). While HIV-1_JRFL_ can be inhibited by small CD4-mimetic compounds, only the most potent are able to do so [10] with (+)-BNM-III-170 having an IC_50_ = 13.49 µM. As we reported previously [18], (*S*)-MCG-IV-210 was unable to inhibit viral infection by HIV-1_JRFL_ (Figure 6), but here we report that several of the derivatives were able to do so. TFH-I-070-A6 presented an IC_50_ of 67.24 µM, TFH-I-116-D1 43.82 µM and ZXC-I-092 54.22 µM (Figure 6B, Table 2). This is the first time that analogs of (*S*)-MCG-IV-210 were able to neutralize HIV-1_JRFL_. We confirmed the potency of these derivative using another tier-2 HIV-1 strain (HIV_AD8,_ Figure 5C, Table 2). TFH-I-116-D1 neutralized HIV-1_AD8_ with an IC_50_ of 4.804 µM, which was similar to that of (+)-BNM-III-170 (IC_50_ = 3.749 µM). Whether the H-bond formation of TFH-I-116-D1 with Env Asp^368^ contributed to the improved potency of this analog remains to be demonstrated. We believe that this derivative poses a first step in the right direction. Additional structural modifications based on the new structural information provided in Figures 4 and 5 are likely to help achieve this goal.

**Figure 6.**
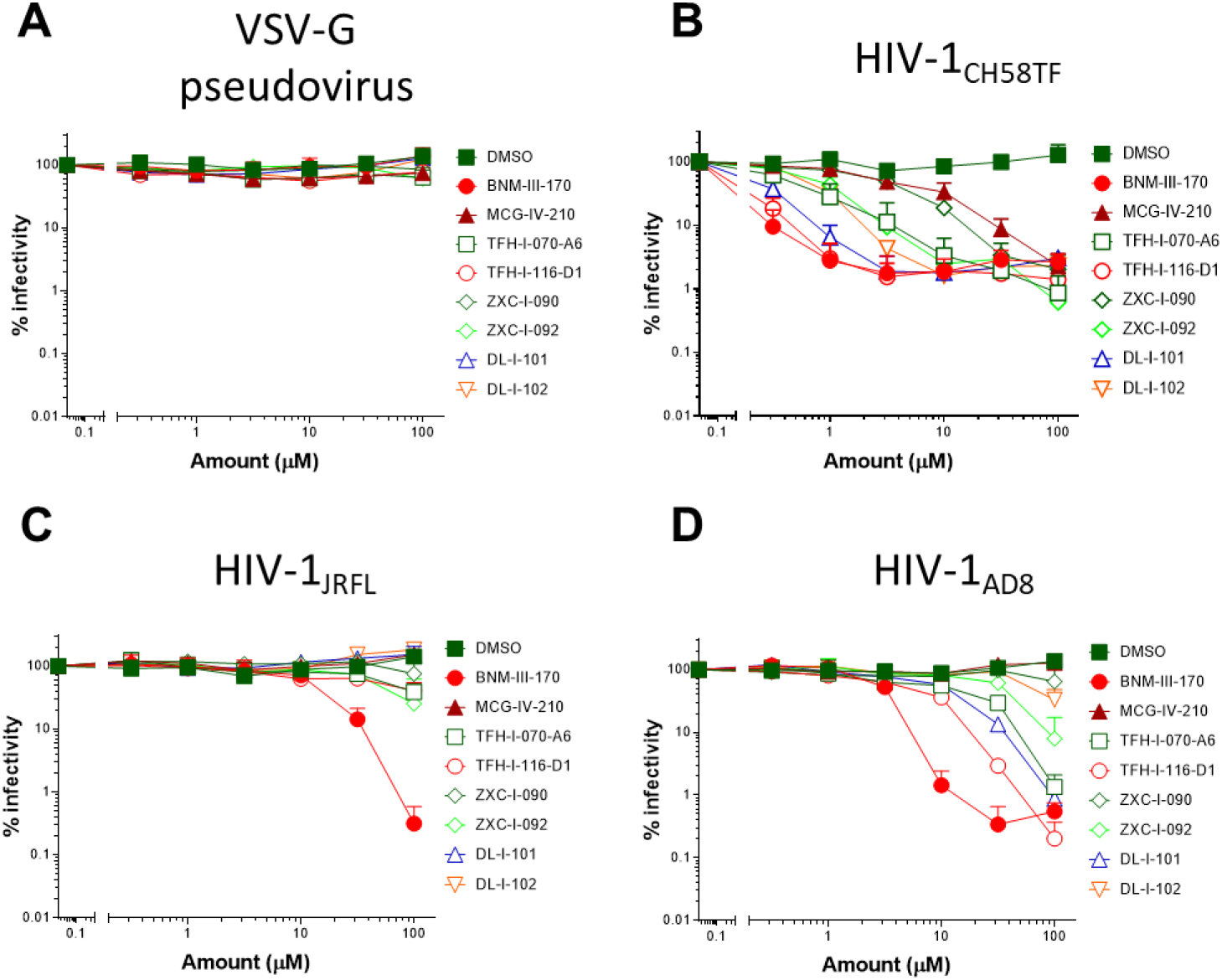
Small CD4-mimetic neutralization of pseudoviral particles. Neutralization of VSV-G pseudovirus (**A**), HIV-1_CH58TF_ (**B**), HIV-1_JRFL_ (**C**) or HIV-1_AD8_ (**D**) virus was done with the indicated amounts of different compounds or the same volume of DMSO, luciferase activity (RLU - relative light units) was measured. Relative infectivity was calculated as the percentage of the value seen in the absence of compound. Data shown are the mean ± SD of at least three independent experiments.

**Table 2.** Viral neutralization. Neutralization of VSV-G HIV-1_CH58TF_, HIV-1_JRFL_ or HIV-1_AD8_ pseudovirus are shown as IC_50_ (average ± SD) of at least three independent experiments; IC_50_ values are in µM.

|  | VSV-G<br>pseudovirus | HIV-1<sub>CH58TF</sub> | HIV-1<sub>JRFL</sub> | HIV-1<sub>AD8</sub> |
| --- | --- | --- | --- | --- |
| DMSO | > 100 | > 100 | > 100 | > 100 |
| BNM-III-170 | > 100 | 0.033 ± 0.01 | 13.49 ± 6.65 | 3.75 ± 1.3 |
| MCG-IV-210 | > 100 | 3.46 ± 1.06 | > 100 | > 100 |
| TFH-I-070-A6 | > 100 | 0.45 ± 0.09 | 67.24 ± 30 | 9.59 ± 3.4 |
| TFH-I-116-D1 | > 100 | 0.065 ± 0.01 | 43.82 ± 16.6 | 4.8 ± 1.7 |
| ZXC-I-090 | > 100 | 2.63 ± 0.5 | > 100 | > 100 |
| ZXC-I-092 | > 100 | 0.75 ± 0.12 | 54.22 ± 22.8 | 34.05 ± 13.8 |
| DL-I-101 | > 100 | 0.16 ± 0.03 | > 100 | 10.21 ± 3.1 |
| DL-I-102 | > 100 | 0.66 ± 0.25 | > 100 | 87.02 ± 39 |

### 2.6. Sensitization of HIV-1-infected cells to ADCC

Since CD4mcs are reported to sensitize HIV-1-infected cells to ADCC mediated by HIV+ plasma [17, 18, 24-29], we next evaluated the susceptibility of primary CD4+ T cells infected with HIV-1_CH58TF_, HIV-1_JRFL_ or HIV-1_AD8_ to ADCC mediated by HIV+ plasma in the absence or presence of (*S*)-MCG-IV-210 derivatives, using a FACS-based assay as previously reported [29, 30]. As expected, the positive control (+)-BNM-III-170 and (S)-MCG-IV-210 enhanced the recognition of HIV-1_CH58TF_ infected cells and their susceptibility to ADCC mediated by HIV+ plasma. This was also the case for the new piperidine CD4mc analogs [TFH-070-A6, TFH-I-116-D1, ZXC-I-090, ZXC-I-092, DL-I-101, DL-I-102] (Figure 7A, 7D). In agreement with a more CD4mc-resistant phenotype observed with HIV-1_JRFL_ infected cells, antibodies from PLWH recognized infected cells only in the presence of TFH-I-116-D1 but not the other derivatives (Figure 7B); this binding was not translated into enhanced ADCC compared to what we observed with (S)-MCG-IV-210 (Figure 7E). We observed a more heterogeneous phenotype with HIV-1_AD8_ infected cells, where (S)-MCG-IV-210 did not promote HIV+ plasma binding to infected cells but TFH-070-A6, TFH-I-116-D1 or DL-I-101 did and translated into ADCC for TFH-I-116-D1 or DL-I-101 (Figure 7C, 7F).

**Figure 7.**
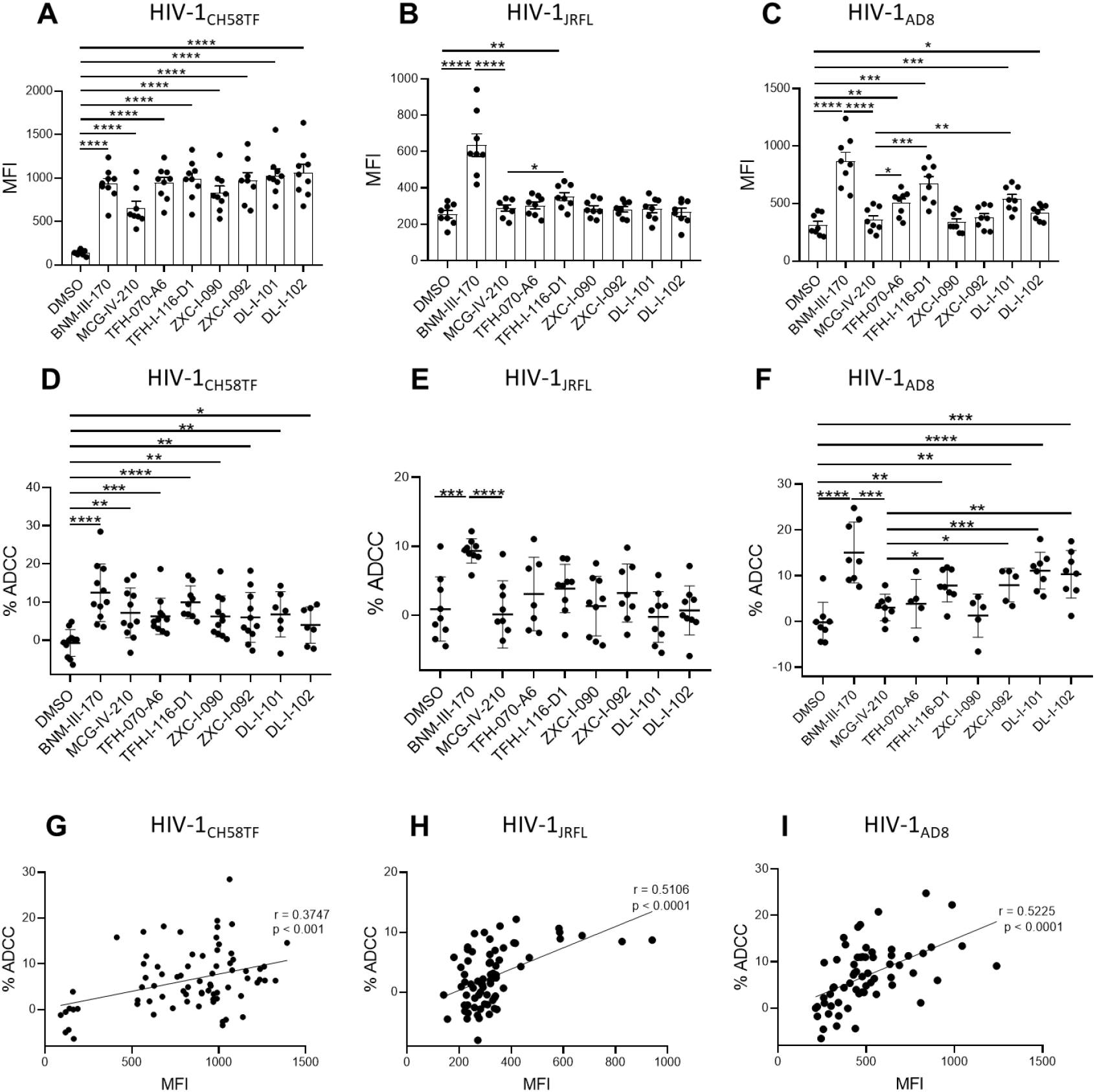
Piperidine CD4mc analogues sensitize HIV-1-infected cells to ADCC. Primary CD4 T cells were infected with HIV-1_CH58TF_ (**A, D**), HIV-1_JRFL_ (**B, E**) or HIV-1_AD8_ (**C, F**). HIV+ plasma (1:1000 diluted) was used for staining (**A** - **C**) or ADCC (**D** – **F**) in the presence of 50µM different compounds or with an equivalent volume of vehicle (DMSO). An Alexa Fluor 647-conjugated anti-human IgG secondary Ab was then used for fluorescent labeling (**A** - **C**). Median fluorescence intensity (MFI) in the presence of compounds or that of DMSO was shown; For ADCC, infected cells were used as target cells in a FACS-based ADCC assay that measures the killing of infected (p24+ cells). The assay determines susceptibility to ADCC mediated by a 1/1,000 dilution of plasma from HIV-1-infected individuals in the presence of 50µM different compounds or with an equivalent volume of vehicle (DMSO). (**G** - **I**) The correlation between cell surface staining with HIV+ plasma and ADCC was calculated using the Pearson r rank correlation. Data shown are the mean ± SD of at least three independent experiments (plasma from more than 5 HIV-1-infected individuals). Statistical significance was evaluated using an unpaired *t* test (*, P < 0.05; **, P < 0.01; ***, P < 0.001; ****, P < 0.0001; ns, not significant).

## 3. Discussion

Small-molecule CD4 mimetics currently attract attention due to their ability both to sensitize HIV-1-infected cells to ADCC [17] and viral particles to neutralization by otherwise ineffective antibodies [31]. *In vivo* studies demonstrated the protection of rhesus macaques from high-dose heterologous transmitted/founder simian-human immunodeficiency virus (SHIV) through the combination of CD4mc (+)-BNM-III-170 and non-neutralizing antibodies elicited by a monomeric gp120 antigen [13]. Moreover, studies in humanized mice supporting NK cell function recently showed that CD4mc in combination with non-neutralizing antibodies decrease viral replication [27, 28] and decrease the level of integrated total DNA in an Fc-dependent manner [28]. To identify additional molecules able to “open-up” Env and expose vulnerable epitopes, we optimized a HTP screening assay using the native trimeric Env to discover a new family of CD4mc able to expose the CoRBS [18]. Further optimization led to (*S*)-MCG-IV-210, a piperidine CD4mc in close proximity with the highly-conserved gp120 Asp^368^ residue that sensitized infected cells to ADCC. Stemming from (*S*)-MCG-IV-210, we explored two structural sites: the C_5_ piperidine ring on the linker and the linear urea, which are important for the binding in the Phe43 cavity and in close proximity to Asp^368^ of the envelope glycoprotein. Screening hundreds of analogs for their ability to expose (a) the CoRBS, to enable recognition of infected cells by HIV+ plasma and (b) inhibit viral infection, followed by structure-based design and evolution, led to six piperidine CD4mc analogs, reported here, with similar or improved potency compared to (*S*)-MCG-IV-210. Structural analysis of this set of analogs, along with that of (+)-BNM-III-170, seems to show that clusters of H-bonds and van der Waals interactions with theβ20-21 are correlated with high potencies, as evidenced by TFH-I-116-D1. This is possibly due to a requirement to stabilize the β20-21 in the CD4-bound-state conformation to promote binding of CoRBS antibodies. Two analogs, DL-I-101 and DL-I-102, donated an H-bond to the α-carboxylic acid group of Asp^368^ and presented improved neutralization and ADCC activities compared to (*S*)-MCG-IV-210, but these compounds displayed less interaction with β20-21 relative to TFH-I-116-D1 and were not superior to it in terms of potency or breadth. This provides a clear direction for further evolution of the piperidine scaffold toward future analogs that simultaneously interact with the highly conserved Asp^368^ and the β20-21 loop, with the goal to improve breadth and potency.

In summary, here we report on the continuing development of piperidine CD4mc. Several new analogs improved the potency to neutralize difficult-to-neutralized HIV-1 strains (JRFL and AD8) and sensitize HIV-1 infected cells to ADCC. Our results grant further development of piperidine CD4mc and explore tighter interactions with gp120 Asp^368^ and the β20-21 loop with the goal to improve potency and breadth.

## 4. Materials and Methods

### 4.1. Ethics Statement

Written informed consent was obtained from all study participants, and research adhered to the ethical guidelines of CRCHUM and was reviewed and approved on October 21 2021 by the CRCHUM institutional review board (ethics committee, approval number CE 16.164 - CA). Research adhered to the standards indicated by the Declaration of Helsinki. All participants were adult and provided informed written consent prior to enrolment in accordance with Institutional Review Board approval.

### 4.2. Cell lines and isolation of primary cells

HEK293T human embryonic kidney cells and TZM-bl cells obtained from ATCC were grown as previously described [7, 17]. Primary human PBMCs, and CD4+ T cells were isolated, activated and cultured as previously described [7, 17]. Briefly, PBMC were obtained by leukapheresis. CD4+ T lymphocytes were then purified from resting PBMCs by negative selection using immunomagnetic beads per the manufacturer’s instructions (StemCell Technologies, Vancouver, BC). CD4+ T lymphocytes were activated with phytohemagglutinin-L (10 µg/mL) for 48 hours and then maintained in RPMI 1640 complete medium supplemented with rIL-2 (100 U/mL).

### 4.3. Chemical synthesis: general considerations

Reactions performed under anhydrous conditions were conducted in oven-dried glassware under an inert atmosphere of argon, unless otherwise stated. Commercial sources of chloroform (ethanol-stabilized), methylene chloride (ethanol-stabilized), toluene, tetrahydrofuran, and diethyl ether were dried over CaH_2_, distilled under reduced pressure, and stored over 4Å molecular sieves under an argon atmosphere. All reagents were purchased from commercial sources and used as received. Reaction mixtures were magnetically stirred under an argon atmosphere, unless otherwise noted, reactions were monitored by either thin-layer chromatography (TLC) with 250-μm SiliaPlate^®^ precoated TLC plates or Waters^®^ ACQUITY analytical ultraperformance liquid chromatography (UPLC) system. Yields refer to chromatographically isolated and spectroscopically pure compounds. Optical rotations were measured on a Jasco P-2000 polarimeter. Proton (^1^H) and carbon (^13^C) nuclear magnetic resonance (NMR) spectra were collected on Bruker Avance III 500 (500 MHz) spectrometer. Chemical shifts (δ) are reported in parts per million (ppm) relative to chloroform-d_3_ (δ 7.26), dimethyl sulfoxide-d_6_ (δ 2.50), acetone-d_6_ (δ 2.05), or methanol-d_4_ (δ 3.31) for ^1^H NMR. Chemical shifts (δ) are reported in parts per million (ppm) relative to chloroform-d_3_ (δ 77.16), dimethyl sulfoxide-d_6_ (δ 39.5), acetone-d_6_ (δ 29.8), or methanol-d_4_ (δ 49.0) for ^13^C NMR. High-resolution mass spectrometry (HRMS) was carried out at the University of Pennsylvania Mass Spectroscopy Service Center on either a (i) Waters LCT Premier XE liquid chromatography-mass spectrometry (LC-MS) system or a (ii) Waters GC-TOF Premier system. Preparative-scale UPLC was performed with a Gilson^®^ SPE Purification system equipped with a Sunfire C_18_ OBD column (10-μm packing material, 30- by 150-mm column dimensions), a 215 liquid handler, a 333 binary gradient module, a 156 UV-visible (UV-Vis) dual-wavelength (254- and 365-nm) detector, and Trilution^®^ 3.0 software. Purification solvent systems were comprised of H_2_O (HPLC-grade) and acetonitrile (HPLC-grade) containing 0.1% trifluoroacetic acid. Supercritical fluid chromatography (SFC) analyses were performed with a Jasco system equipped with a PU-280-CO_2_ plus CO_2_ delivery system, a CO-2060 plus intelligent column thermostat/selector, an HC-2068-01 heater controller, a BP-2080 plus automatic back pressure regulator, an MD-2018 plus photodiode array detector (200 to 648 nm), and PU 2080 plus intelligent HPLC pumps. The purity of new compounds was evaluated by NMR and UPLC-MS (>95%).

### 4.4. Viral production and infection of primary CD4+ T cells

HIV-1 viruses were produced and titrated as previously described [6, 18]. Briefly, plasmids expressing the following full-length infectious molecular clones (IMCs) of HIV-1_CH58TF_, HIV-1_JRFL_ or HIV-1_AD8_ were transfected in 293T cells by standard calcium phosphate transfection. Two days after transfection, cell supernatants were harvested, clarified by low-speed centrifugation (5 min at 1,500 rpm), and concentrated by ultracentrifugation for 1 h at 4°C at 143,260 × g over a 20% sucrose cushion. Pellets were harvested in fresh RPMI, and aliquots were stored at −80°C until use. Viruses were then used to infect activated primary CD4+ T cells from healthy HIV-1 negative donors by spin infection at 800 × *g* for 1 h in 96-well plates at 25 °C.

### 4.5. Viral neutralization

The viral infection assay was done as previously described [32]. Briefly, TZM-bl target cells were seeded at a density of 5 × 10^3^ cells/well in 96-well luminometer-compatible tissue culture plates (Perkin Elmer) 24 h before infection. HIV-1_CH58TF_, HIV-1_JRFL_ or HIV-1_AD8_ viruses in a final volume of 100 µl was incubated with indicated amount of different compounds or the same volume of DMSO for one hour at 37°C, then the mixture was added to the target cells followed by incubation for 48 h at 37°C; the medium was then removed from each well, and the cells were lysed by the addition of 30 µl of passive lysis buffer (Promega) and three freeze-thaw cycles. A LB 941 TriStar luminometer (Berthold Technologies) was used to measure the luciferase activity of each well after the addition of 100 µl of luciferin buffer (15 mM MgSO_4_, 15 mM KPO_4_ [pH 7.8], 1 mM ATP, and 1 mM dithiothreitol) and 50 µl of 1 mM D-luciferin potassium salt (Prolume).

### 4.6. Antibodies and plasma

The anti-CoRBS 17b mAb was used alone or in combination with different compounds for cell-surface staining. Plasma from different HIV-infected donors were collected, heat-inactivated and conserved as previously described [7, 17]. Alexa Fluor 647 conjugated Goat anti-human antibodies (Invitrogen) were used as secondary Abs.

### 4.7. Flow cytometry analysis of cell-surface staining

Cell-surface staining was performed as previously described [15-17]. Primary CD4 T cells were isolated from healthy donors and infected with HIV-1_CH58TF_, HIV-1_JRFL_ or HIV-1_AD8_. Binding of HIV-1-infected cells by plasma (1:1,000 dilution) or 17b mAb (5μg/ml) in the presence or absence of 50 μM compounds was performed 48 hours after infection. Cells were then incubated at 37 °C for 1 hour followed by adding anti-human Alexa Fluor-647 (Invitrogen) secondary Abs for 20 minutes. Cells were then stained intracellularly for HIV-1 p24, using the Cytofix/Cytoperm Fixation/ Permeabilization Kit (BD Biosciences, Mississauga, ON, Canada) and the fluorescent anti-p24 mAb (PE-conjugated anti-p24, clone KC57; Beckman Coulter/Immunotech). The percentage of infected cells (p24+ cells) was determined by gating the live cell population on the basis of the AquaVivid viability dye staining. Samples were analyzed on an LSRII cytometer (BD Biosciences), and data analysis was performed using FlowJo vX.0.7 (Tree Star, Ashland, OR, USA).

### 4.8. ADCC FACS-based assay

Measurement of ADCC using the FACS-based assay was performed at 48h post-infection as previously described [7, 17, 33, 34]. Briefly, HIV-1_CH58TF_, HIV-1_JRFL_ or HIV-1_AD8_ infected primary CD4+ T cells were stained with viability (AquaVivid; Thermo Fisher Scientific) and cellular (cell proliferation dye eFluor670; eBioscience) markers and used as target cells. Autologous PBMC effectors cells, stained with another cellular marker (cell proliferation dye eFluor450; eBioscience), were added at an effector: target ratio of 10:1 in 96-well V-bottom plates (Corning, Corning, NY). Then the mixed cells were incubated with HIV+ plasma (1:1000), in the presence of 50 µM of compounds or with equivalent volume of vehicle (DMSO). The plates were subsequently centrifuged for 1 min at 300 g, and incubated at 37°C, 5% CO2 for 4 to 6 h before being fixed in a 2% PBS-formaldehyde solution. Samples were analyzed on an LSRII cytometer (BD Biosciences). Data analysis was performed using FlowJo vX.0.7 (Tree Star). The percentage of ADCC was calculated with the following formula: (% of p24+ cells in Targets plus Effectors) − (% of p24+ cells in Targets plus Effectors plus plasma) / (% of p24+ cells in Targets) by gating on infected lived target cells.

### 4.9. Statistical analysis

Statistics were analyzed using GraphPad Prism version 7.0a (GraphPad, San Diego, CA, USA). Every data set was tested for statistical normality and this information was used to apply the appropriate (parametric or nonparametric) statistical test. P values <0.05 were considered significant; significance values are indicated as * p<0.05, ** p<0.01, *** p<0.001, **** p<0.0001.

### 4.10. CRF01_AE core e expression and purification

Clade AE LM/HT and LM/HS 93TH057gp120_coree_ protein was produced by transfection into GnTI-293F cells. Cells were grown in suspension for 7 days at 37° C and 90% humidity. The cells were pelleted by centrifugation and the medium was filtered through a 0.2 micron filter. Protein was purified on a 17b affinity column (17b IgG covalently linked to protein A agarose) equilibrated with phosphate buffered saline (PBS) pH 7.2. The column was washed with PBS and gp120 eluted with 0.1 M glycine pH 3.0. Eluted fractions were immediately diluted 10:1 with 1 M tris(hydroxymethyl)aminomethane-HCl (Tris-HCl) pH 8.5. Eluted protein was concentrated to approximately 10 mg/ml and the buffer was then exchanged to 50 mM sodium acetate pH 6.0 and 350 mM sodium chloride. EndoH_f_ (New England Biolabs) was added and the sample was equilibrated overnight at 37° C to deglycosylate the protein. Deglycosylated protein was then passed over an amylose column equilibrated in 25 mM Tris-HCl pH 7.2 and 200 mM sodium chloride to remove EndoH_f_ (Maltose binding protein tagged EndoH). The protein was concentrated and the sample loaded onto a Superdex 200 gel filtration column (Cytiva) equilibrated in 10 mM Tris-HCl pH 7.2 and 100 mM ammonium acetate. Fractions corresponding to the deglycosylated gp120 size were concentrated to approximately 10 mg/ml for use in crystallization trials.

### 4.11. Crystallization of gp120 cores complex with CD4mc

Crystals were grown with the hanging drop method from 10% polyethylene glycol (PEG) 3350, 5% PEG 400, and 0.1 M 4-(2-hydroxyethyl)-2-piperazineethanesulfonic acid (HEPES) pH 7.5. Crystals usually appeared within 1 to 2 weeks when incubated at 21° C. CD4 mimetic compounds were added by soaking crystals in 1 mM of the compound. Crystals were then frozen for data collection. Prior to freezing crystals were briefly soaked in the crystallization condition and compound supplemented with 20% of 2-Methyl-2,4-pentanediol (MPD) as a cryoprotectant.

### 4.12. Data Collection, Structure Solution and Refinement

Diffraction data were collected at the Stanford Synchrotron Radiation Light Source (SSRL) beamlines 9-2 and 12-2 on a Dectris Pilatus 6M area detector. All data were processed and reduced with HKL3000 [35] or MOSFLM and SCALA from the CCP4 suite [36]. Structures were solved by molecular replacement with PHASER from the CCP4 suite [36] based upon the coordinates from PDB ID 6ONF. Refinement was carried out with Refmac5 [36] and/or Phenix [37] and model building was done with COOT [36]. Data collection and refinement statistics are shown in **Table 1**. Ramachandran statistics were calculated with MolProbity and illustrations were prepared with Pymol Molecular graphics (http://pymol.org).

## Author Contributions

S.D., M.P., A.B.S and A.F. conceived the study. S.D., M.P., A.B.S. and A.F. designed experimental approaches. S.D., W.D.T., H.Z., D.L., T.H., X.Z., D.N., R.S., D.V., J.R., G.G.L., H.M., M.M., and A.F. performed the syntheses, analyzed, and interpreted the experiments. S.D. performed statistical analysis. S.D., W.D.T., H.Z., D.L., C.A., A.B.S., M. P. and A.F. wrote the manuscript with inputs from others. Every author has read, edited and approved the final manuscript.

## Disclaimer

The views expressed in this manuscript are those of the authors and do not reflect the official policy or position of the Uniformed Services University, the U.S. Army, the Department of Defense, the National Institutes of Health, the Department of Health and Human Services or the U.S. Government, nor does mention of trade names, commercial products, or organizations imply endorsement by the U.S. Government.

## Institutional Review Board Statement

The study was approved by the CRCHUM institutional review board (ethics committee, approval number CE 16.164 - CA)

## Informed Consent Statement

Informed consent was obtained from all subjects involved in the study.

## Data Availability Statement

Data is contained within the article and supplementary materials.

## Acknowledgments

The authors are grateful to the donors who participated in this study. The authors thank the CRCHUM BSL3 and Flow Cytometry Platforms for technical assistance, and Mario Legault from the FRQS AIDS and Infectious Diseases network for cohort coordination and clinical samples. Use of the Stanford Synchrotron Radiation Lightsource, SLAC National Accelerator Laboratory, is supported by the U.S. Department of Energy, Office of Science, Office of Basic Energy Sciences under Contract No. DE-AC02-76SF00515. The SSRL Structural Molecular Biology Program is supported by the DOE Office of Biological and Environmental Research, and by the National Institutes of Health, National Institute of General Medical Sciences. The funders had no role in study design, data collection and analysis, decision to publish, or preparation of the manuscript and the contents of this publication are solely the responsibility of the authors.

